# Extended amygdala corticotropin-releasing hormone neurons regulate sexually dimorphic changes in pair bond formation following social defeat in prairie voles (*Microtus ochrogaster*)

**DOI:** 10.1101/2024.11.11.623026

**Authors:** Maria C. Tickerhoof, Lina K. Nerio Morales, Jeff Goff, Erika M. Vitale, Adam S. Smith

## Abstract

The neurobiological mechanisms underlying the connection between anxiety brought on by social stressors and the negative impact on relationship formation have remained elusive. In order to address this question, we used the social defeat model in the socially monogamous prairie vole to investigate the impact of this stress on pair bond formation. Social defeat experience inhibited partner preference formation in males but promoted preference in females. Furthermore, pair bonding increased corticotropin-releasing hormone (CRH) expression in the bed nucleus of the stria terminalis (BNST) in male prairie voles, while defeat experience increased BNST CRH expression in females. Chemogenetic excitation of BNST CRH neurons during a short cohabitation with a new partner promoted a partner preference in stress-naïve prairie voles. Interestingly, chemogenetic inhibition of BNST CRH neurons during cohabitation with a new partner blocked partner preference in stress-naïve males but promoted preference in defeated males. Inhibition of BNST CRH neurons also blocked partner preference in stress-naïve females but did not alter preference behavior in defeated females. This study revealed sexual dimorphism in not only the impact of social defeat on pair bond formation, but also in the role BNST CRHergic neurons play in regulating changes in pair bonding following social conflict.

## Introduction

Approximately one in three individuals in the United States are affected by social anxiety during their lifetime [1,2]. The prevalence of social anxiety disorder is 27% higher in individuals that experience social conflict, such as bullying and other forms of peer victimization [3–5]. The fear and avoidance of social interactions demonstrated by those with high social anxiety and social anxiety disorder contributes to reduced social network size and considerable difficulty in forming new social attachments, including romantic relationships [6–9]. Thus, social anxiety may lead to limited access to the multiple mental and physical health benefits of close relationships [10–12], implying long-lasting negative consequences for the well-being of those that have experienced social conflict beyond the initial acute stress. Despite this established connection between social anxiety and difficulties with social attachment, the underlying neurobiological mechanisms have received limited investigation.

The bed nucleus of the stria terminalis (BNST) is a region that could contribute to the connection between social conflict and attachment. The BNST is widely acknowledged for its role in stress, anxiety, and responses to unpredictable threats [13–18]. However, in mice, both aversive and appetitive behavior are regulated by distinct neuronal populations within the BNST [19]. Beyond this, the BNST may also be involved in social attachment as it promotes maternal care and other prosocial behavior in rats and mice, as well as pair bonding between adult mating pairs in prairie voles (*Microtus ochrogaster*) [20–24]. The BNST is also highly reactive to opposite-sex social exposure and is a notable regulator of sexually-motivated social behavior in multiple rodent species [25–28]. This intersection of roles in both social behavior and threat assessment leads to the BNST being key to phenomena like social vigilance following social conflict in California mice (*Peromyscus californicus*) [29] and opens the possibility that this region has distinct influences on social attachment during periods of high social anxiety.

Corticotropin-releasing hormone (CRH) is a neuropeptide with critical roles in multiple biobehavioral systems. Extra-hypothalamic CRH, including CRH originating from the BNST, is known to regulate the behavioral response to various stressors in mice, including social conflict [30–32]. However, CRH has been repeatedly demonstrated to regulate behavioral responses not only to stress and aggression, but also rewarding stimuli such as social and sexual interaction in multiple rodent species [33–37]. Notably, CRH production within the BNST is increased as a result of social attachment [38]. Because of these dual roles of BNST CRH production and activity in both anxiety- and affiliation-associated contexts, it is likely that stress-sensitive CRH-producing neuronal populations (hereafter referred to as “CRHergic”) play a role in the aforementioned differential regulation of both appetitive and aversive behavior by the BNST [19]. Thus, CRHergic neurons in the BNST could underlie the connection between social conflict and reduced social affiliation.

In order to investigate this hypothetical role of BNST CRHergic neurons, an animal model capable of modeling both social conflict and social attachment is necessary. For this reason, this study made use of the social defeat model in the socially monogamous prairie vole [39–42]. Male-female mating pairs of this species form long-lasting and robust pair bonds which have been used to study the neurobiological underpinnings of social attachment [43], including the impact of experiences like chronic stress on pair bond formation [44]. However, the impact of social stress on pair bond formation has yet to be investigated. Thus, we aimed to characterize the impact of social defeat on pair bonding in both sexes and the role of BNST CRHergic neurons in regulating pair bond formation both before and following social conflict. Due to the negative impact of chronic stress on pair bonding in males [44] and the social avoidance state induced by social defeat in prairie voles of both sexes [39,40], we hypothesized that social defeat would lead to impairments in pair bonding in both males and females. Additionally, we hypothesized that BNST CRHergic neurons would be a novel neurobiological mechanism promoting pair bond formation in a stress-naïve state and would contribute to any observed effects of social defeat on bonding.

## Materials and Methods

### Animals

Subjects were captive-bred sexually naïve male and female prairie voles descended from populations captured in southern Illinois. Voles were weaned on postnatal day 21 and housed with a same-sex non-littermate in cages (29.2 L x 19.1 W x 12.7 H cm) containing corn cob bedding and crinkle paper nesting material with food and water *ad libitum*. Except during periods where animals were being conditioned or during behavioral testing, animals were housed in these conditions with the same animal they had been housed with since weaning (designated as the “home cage”). At the time of pairing, subjects were paired with an opposite-sex conspecific in similar housing conditions as those in which they had lived in since weaning. Colony rooms were maintained on a 14L:10D photoperiod (lights on at 0600 hr) and at a temperature range of 21 ± 1°C. Subjects were 80-110 days old and 25-50 grams in weight at the initiation of their experimental timeline. All procedures were conducted in accordance with the National Institutes of Health Guide for the Care and Use of Laboratory Animals and the Institutional Animal Care and Use Committee at the University of Kansas.

## Behavioral paradigms

### Social defeat

Social defeat was performed as previously described [39,40,42]. Briefly, on the days of defeat sessions, subjects and resident aggressors were brought up to the behavior testing room and left to habituate for at least 30 minutes. Resident aggressor pairs were separated so that the resident that was the same sex as the subject was left in the home cage. The subjects were introduced into the resident home cage and monitored for physical aggression. The resident aggressor and subject were separated upon reaching seven physical attacks or 15 minutes, whichever came first. The subject was then placed in a corral in the center of the home cage for the remainder of an hour, then returned to their home cage. Defeat was repeated for three days with a different resident each day. Non-defeated control animals were separated from their cage mate and placed in a novel empty cage for an hour, then returned to the home cage. This protocol reliably induces a social avoidance phenotype in both male and female prairie voles [39,40,42].

### Social preference/avoidance test

Social interaction was characterized with a social preference/avoidance test (SPA) as previously described [39,40,42], with the exception that the social stimulus was the opposite sex as the subject. The subject was introduced into the center chamber of a 75 L x 20 W x 25 H cm three-chamber arena and left to habituate to the testing arena for ten minutes. After the ten-minute habituation, the subject was sequestered into the center chamber and the test stimuli (rubber water bottle stopper for *object*, unknown opposite-sex conspecific for *social*; both in metal corrals) were placed against opposite walls of the side chambers. The locations of object/social stimuli were counter-balanced in a pseudorandom manner. The barriers were removed and the subject was left to explore and interact with each stimulus for ten minutes. Olfactory investigation (sniffing) was quantified throughout the ten-minute interaction period. A significant preference for the social stimulus over the object stimulus is typical behavior for prairie voles, while a lack of preference between the two stimuli is indicative of a social avoidance phenotype [39,40,42].

### Partner preference test

Partner affiliation was characterized with partner preference test (PPT) as previously described [45–47]. The subject’s partner and an unknown conspecific the opposite sex from the subject (stranger) were tethered in opposite chambers of a 75 L x 20 W x 25 H cm three-chamber arena. The locations of partner/stranger stimuli were counter-balanced in a pseudorandom manner. The subject was placed in the center chamber separated by barriers. After lifting the barriers, subjects were left to explore the arena and interact with each animal for three hours. Stationary contact with the stimuli was quantified as a measure for partner preference. Preferential stationary contact with the partner over the stranger (partner preference) is the standard measurement indicative of pair bond formation in prairie voles [47].

#### qRT-PCR

Fresh brains flash frozen on dry ice, then stored at -80° C until sectioning. The brains were sectioned at 300 μm and the BNST was punched out with a 1 mm biopsy punch. Tissue punches were homogenized, and mRNA was extracted and purified using an RNEasy mini kit (Qiagen Cat# 74104). RNA was quantified using a Qubit RNA High Sensitivity kit (Invitrogen Cat# Q32852). RT-PCR was performed as described according to a standard protocol [48,49]. mRNA was converted to cDNA using a high-capacity cDNA reverse transcription kit (Applied Biosystems Cat# 4368814). The ΔΔCT method was used to quantify fold change between groups with HPRT acting as a control gene [50].

## Chemogenetic manipulation and validation

### Drug preparation

Colchicine (Thermo Scientific Cat# J61072ME) was dissolved in ddH_2_O at a concentration of 60 mg/mL, then aliquoted and stored at −20°C until the day of use. On the day of injection, one aliquot was thawed and diluted with sterile 0.9% saline to a concentration of 20 mg/mL.

Clozapine-N-oxide (CNO) was obtained from the NIMH Chemical Synthesis and Drug Supply Program. CNO was dissolved in DMSO at a concentration of 25 mg/mL, diluted with ddH_2_O to a final concentration of 3.45mg/mL, then aliquoted and stored at −20° C until the day of use. On the day of use, aliquots were thawed and diluted with 0.9% sterile saline to a final concentration of 0.9 mg/mL.

### Viral vector preparation

The viruses used were as follows: AAV2-CRH-Cre (Biohippo Cat# PT-0588), AAV2-hSyn-DIO-mCherry (mCherry; Addgene Cat# 50459-AAV2), AAV2-hSyn-DIO-hM3D(Gq)-mCherry (Gq; Addgene Cat# 44361-AAV2), and AAV2-hSyn-DIO-hM4D(Gi)-mCherry (Gi; Addgene Cat# 44362). Viruses were aliquoted upon receipt and stored at −80o C until day of use. On the day of injection, one aliquot each of the CRH-Cre virus and appropriate Cre-inducible virus were thawed and combined in a 50/50 ratio prior to loading.

### Stereotactic injections

Animals were anesthetized with a ketamine/dexmedetomidine cocktail (75/1 mg/kg, intraperitoneal) and headmounted into a stereotax (Stoelting). For viral injections, the viral vector cocktail was bilaterally injected via syringe (Hamilton Cat# 65458-02) into the anterodorsal BNST (A/P +0.30, M/L ±0.95, D/V −4.95 relative to bregma) at 75 nL/min to a final volume of 500nL per hemisphere. The needle was left in place for a 5-minute diffusion period before removing. For colchicine injection, 20mg/mL colchicine in 0.9% saline was unilaterally injected into the lateral ventricle (A/P +0.30, M/L - 0.95, D/V −2.5) at 250nL/min to a total final volume of 1μL (20μg). Anesthesia was reversed with atipamezole (2.5 mg/kg, intraperitoneal) and animals were returned to a clean cage for recovery.

### Perfusions

Animals were anesthetized with a ketamine/dexmedetomidine cocktail (75/1 mg/kg), exsanguinated with 0.9% saline, then perfused with 4% paraformaldehyde in 0.1M phosphate buffer. Brains were extracted and post-fixed in 4% paraformaldehyde in 0.1M phosphate buffer for 2-3 hours then transferred to 30% sucrose in 0.1M phosphate buffer for 2-3 days. Brains were then flash frozen on crushed dry ice and stored at −80° C until sectioning.

### Validation of viral distribution

For all perfused brains other than those injected with colchicine, brains were cryosectioned at 30μm and mounted onto a glass slide. Brains injected with colchicine were cryosectioned at 30μm and placed free-floating in 0.1M PBS with 0.1% sodium azide before processing for CRH-ir as described below. Slides were coverslipped with in-house prepared gelvatol and imaged on a light microscope (Leica). mCherry expression regional specificity was characterized using the Paxinos and Watson Mouse Brain Atlas as a guide for anatomical markers. Animals were removed if viral mCherry expression was observed outside of the BNST.

### CRH immunoreactivity

Perfused brains were sectioned at 30 μm to collect the BNST and stored in PBS with 0.1% sodium azide at 4° C until immunolabeling. On the day of immunolabeling, sections were brought to room temperature then rinsed 3 x 5 min in PBS, then 1 x 10 min in 1% NaBH_4_ in PBS, then finally 3 x 5 min in PBS. After rinsing, sections were incubated in 10% normal goat serum (NGS) in T-PBS (PBS with 0.5% Triton X-100) for 1 hour. Sections were then incubated in anti-CRH primary antibody in T-PBS with 2% NGS for 48 hours at 4° C. After 48h at 4° C, sections continued to be incubated in primary antibody for 1 hour at room temperature. Sections were then rinsed in T-PBS 3 x 5 min, followed by incubation in secondary antibody for 2 hours at room temperature. Sections were then rinsed again 1 x 5 min in T-PBS, incubated in DAPI in PBS 1 x 5 min, rinsed in PBS 2 x 5 min, then finally mounted onto glass microscope slides and cover slipped using gelvatol for imaging.

## Experimental Design

### Effect of social defeat on social approach-avoidance behavior and pair bond formation

Male and female prairie voles were separated into Control and Defeat conditions (detailed in *Behavioral paradigms*). One week after the final day of conditioning, subjects had sociality assessed in the three-chamber SPA test with a novel opposite-sex animal (detailed in *Behavioral paradigms*). Immediately after testing, subjects were paired with the same opposite-sex animal and left to cohabitate for 24 hours (detailed in *Behavioral paradigms*). Males in the 24-hour cohabitation conditions had female partners treated with subcutaneous estradiol benzoate (EB) for three days prior to pairing to promote mating, as mating increases the chance of pair bond formation in males but is not necessary for females [46,51,52]. Cohabitation behavior was recorded throughout the entire cohabitation to confirm whether or not mating had occurred; mating was confirmed to have occurred in all pairings where the females were administered EB, and no mating occurred in the pairings without EB treatment. Immediately at the end of cohabitation, pair bond formation was assessed with PPT (detailed in *Behavioral paradigms*). The experiment was repeated with a separate cohort of females with only 6 hours of cohabitation, which is typically deemed insufficient to lead to pair bond formation [52], cf. [53].

### Effect of social defeat and pair bonding on BNST CRH expression

Male and female prairie voles were separated into three groups: sexually-naïve, 6-hour cohabitation, and 24-hour cohabitation. Animals in the sexually-naïve group remained with their cage mates, and subjects in the other two groups were paired with an opposite-sex partner and left to cohabitate for 6 or 24 hours. At the end of the cohabitation period animals were sacrificed via rapid decapitation and brains were harvested and processed as described.

### Validation of CRH-Cre selective expression

12 days following injection of CRH-Cre and mCherry into the BNST, 20 μg colchicine was injected unilaterally into the lateral ventricle. Since CRH is difficult to label in the BNST, colchicine is used to assist with increasing accumulation of CRH within the cell body and decrease fiber labeling [54]. Although colchicine administration has been demonstrated to lead to accumulation of neuropeptides in cells that may not normally synthesize such compounds under physiological conditions [55], it is appropriate to use in the context of validating the selective transcription of viral vectors in cells that can produce CRH. Two days after the colchicine injection animals were perfused and brains harvested, then tissue was processed for CRH immunoreactivity to quantify colocalization of the viral products in CRH neurons. 2-3 sections from a cohort of 6 animals injected with the CRH-Cre/DIO-mCherry followed by colchicine were used for viral expression validation. Cells expressing red label alone (CRH-Cre+, CRH-) and red label plus green label (CRH-Cre+, CRH+) were quantified. The ratio of red+green over (red+green)+(red alone) was used to determine the selectivity of viral expression to CRH cells.

### Activation of BNST CRHergic neurons in stress-naïve animals

Male and female prairie voles received a bilateral injection into the BNST of CRH-Cre with either mCherry or Gq. Two weeks after surgery, animals were administered an intraperitoneal injection of 3 mg/kg CNO. Half an hour later, animals were paired with an opposite-sex mate and left to cohabitate for 6 hours. This period of time without mating has been demonstrated to be insufficient to allow for formation of partner preference in both males and females [46,51,52,56]. Female partners of male subjects were not EB-primed for this experiment, and cohabitation behavior was recorded and later observed to confirm the absence of mating in all pairs. Partner preference behavior was characterized immediately at the end of cohabitation. Animals were perfused and brains harvested the following day for validation of viral expression.

### Inhibition of BNST CRHergic neurons in stress-naïve animals

Male and female prairie voles received a bilateral injection into the BNST of CRH-Cre with either mCherry or Gi. Four weeks after surgery, animals were administered an intraperitoneal injection of 3 mg/kg CNO. Half an hour later, subjects were paired with an opposite-sex conspecific and left to cohabitate for 24 hours. Female partners of male subjects were treated with EB to promote mating, and cohabitation behavior was recorded and later observed to confirm the presence of mating for all male subjects and absence of mating for all female subjects. PPT was performed immediately at the end of cohabitation. Animals were perfused and brains harvested the following day for validation of viral expression.

### Inhibition of BNST CRHergic neurons in defeated animals

Experimental design was identical to the section above, with the following two exceptions: 1) 10 days prior to pairing, all subjects underwent the three-day social defeat conditioning protocol; 2) male subjects were paired with a female partner for 24 hours, and female subjects were paired with a male partner for 6 hours. Female partners of male subjects were treated with EB to promote mating, and cohabitation behavior was recorded and later observed to confirm the presence of mating for all male subjects and absence of mating for all female subjects.

## Quantification and statistical analysis

Prior to analysis, any animals in Gq and Gi groups with expression outside the BNST were excluded from the dataset. Behavioral analysis was performed by manual scoring using JWatcher software (UCLA) by an observer blind to experimental condition. Viral validation of selective expression was quantified via manual counting of cells using ImageJ software. Statistical analysis was performed using SPSS (IBM). Male and female RT-PCR data were analyzed by two-way ANOVA (pairing x defeat). Partner preference behavior was analyzed by paired-sample t-test (partner vs. stranger). For SPA, social interaction/avoidance was analyzed by mixed model ANOVA (between subject: sex x condition; within subject: stimulus type).

## Results

### BNST CRH expression increases in defeated males and pair bonded females

There were main effects of social defeat experience on BNST CRH mRNA expression in male prairie voles (n = 6-8 per group; F_1,23_ = 9.499, p = 0.005; *Figure 1A*) and male cohabitation on BNST CRH mRNA expression in female prairie voles (n = 6-8 per group; F_1.23_ = 13.139, p = 0.001; *Figure 1B*). Specifically, defeated males had significantly higher CRH mRNA expression in the BNST compared to stress-naïve males. Further, females living with a male partner for 24 hours had significantly greater expression of BNST CRH mRNA than sexually-naïve females. There were no other main effects or interactions for males or females.

**Figure 1:**
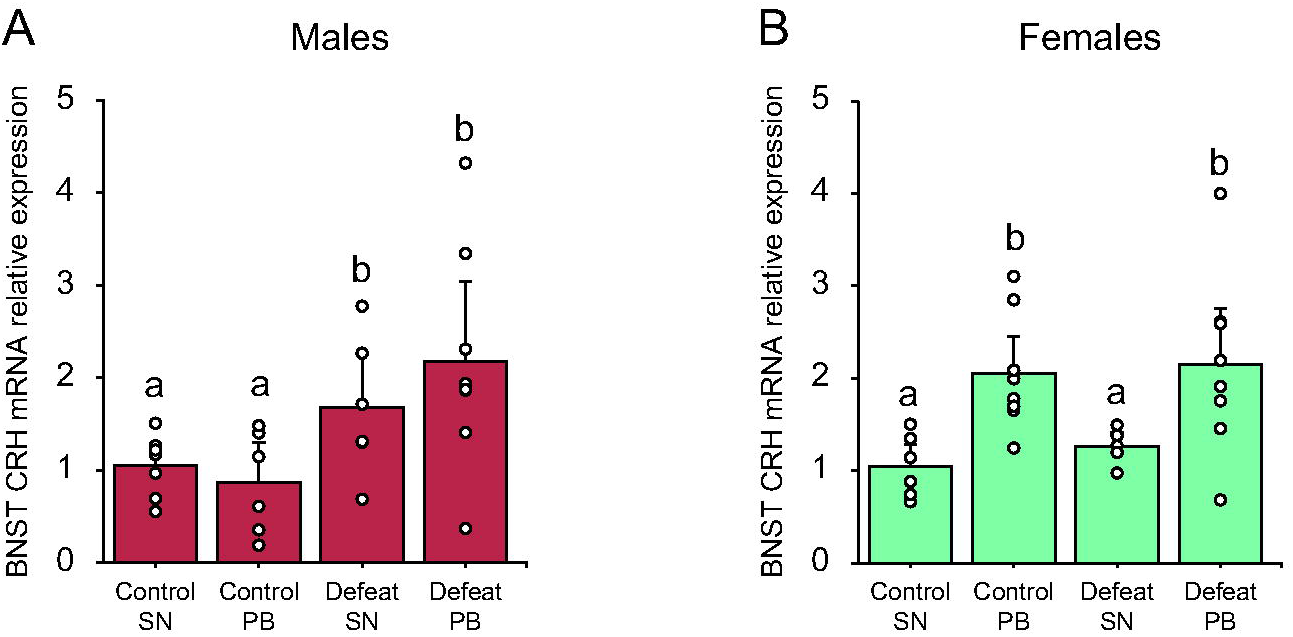
Effects of cohabitation with an opposite-sex partner with or with defeat experience on BNST CRH mRNA expression. A) BNST CRH mRNA expression was increased by social defeat experience but was unaffected by cohabitation with a female conspecific in male prairie voles (n = 7-8 group). B) By contrast, social defeat experience did not affect BNST CRH mRNA expression in female prairie voles but cohabitation with a male conspecific increased it (n = 7-8 group). Different letters above each bar denotes statistical differences between groups, p<0.05. SN = sexually-naïve, PB = pair bonded.

### Social defeat leads to sexually dimorphic changes in pair bond formation

We initially hypothesized that social defeat would impair pair bond formation in both male and female prairie voles. Male and female prairie voles were subjected to either control or defeat conditioning, then a week later tested for social investigation of a potential mate with the SPA test. Immediately following SPA, subjects were paired with the opposite-sex social stimulus for a 24-hour cohabitation, then pair bonding tested with the PPT (*Figure 2A*). A 24-hour cohabitation time was selected as it reliably leads to expression of partner preference behavior (a marker for the successful formation of a pair bond) in both sexes under stress-naïve conditions [46,51,52].

**Figure 2:**
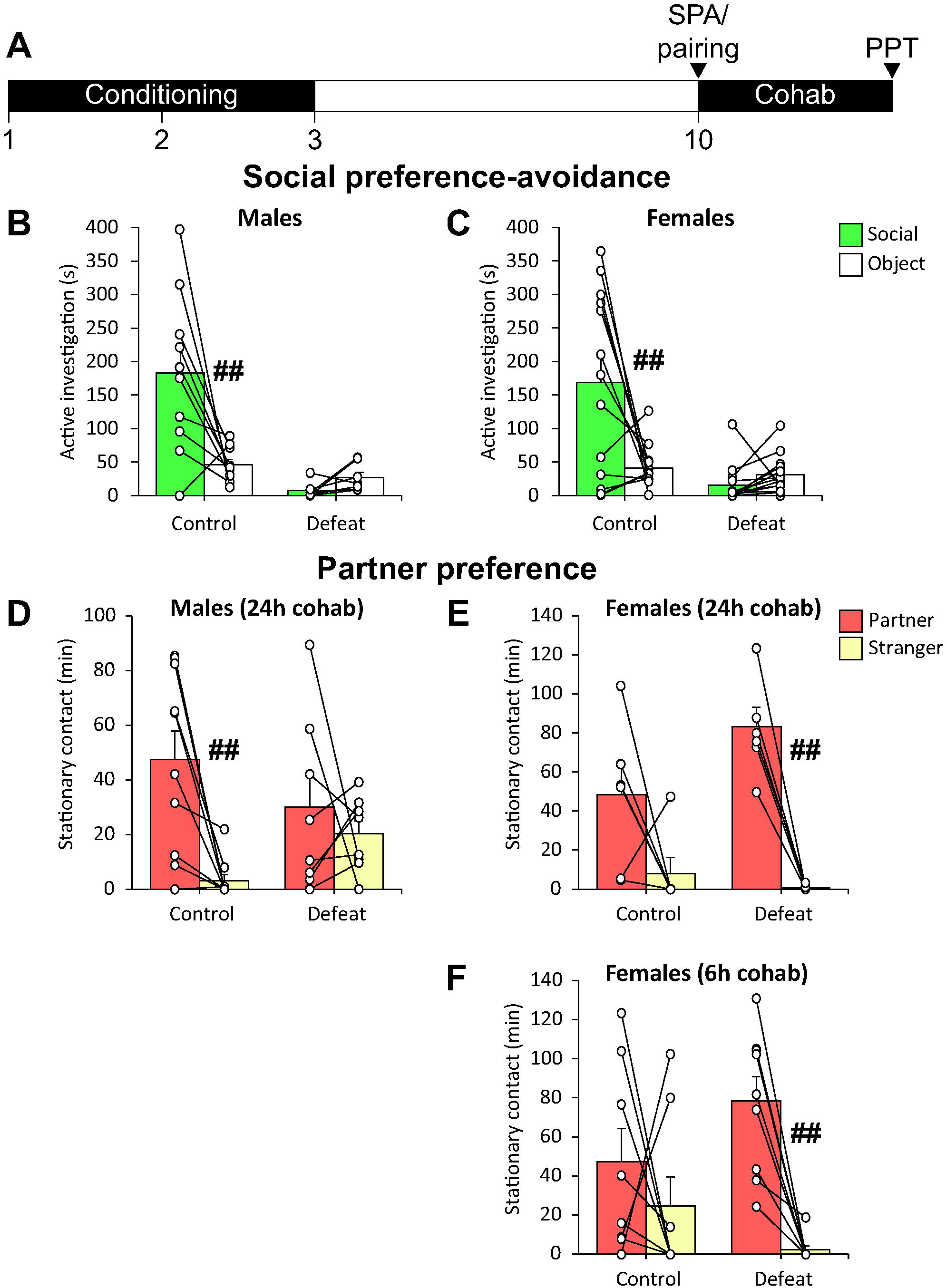
Impact of social defeat on opposite-sex social investigation and pair bond formation. A) Experimental timeline. Subjects underwent three days of control or defeat conditioning, then were paired with an opposite-sex partner immediately following SPA testing. Partner preference was evaluated at the end of a 6- or 24-hour cohabitation. Social defeat led to social avoidance of an opposite-sex conspecific in both B) males (n = 10 control/7 defeat) C) and females (n = 13 control/14 defeat). Social defeat also caused an impairment in pair bond formation D) in males following 24 hours of cohabitation (n = 10 control/8 defeat), E) but not in females following 24 hours of cohabitation (n = 6 control/6 defeat). F) Social defeat led to an acceleration in pair bond formation in females following 6 hours of cohabitation (n = 8 control/9 defeat). ##p < 0.01 vs. other stimulus.

Prior to pairing, the SPA test revealed that social defeat experience eliminated social preference of a novel opposite-sex conspecific over a novel object in both males and females (n = 7-14 per group; F_1,40_ = 20.312, p = 0.00006; *Figure 2B-C*). Following the 24-hour cohabitation, control males demonstrated a significant preference for their partner over the stranger (n = 10; t_9_ = 4.064, p = 0.003), but defeated males had no preference between the partner and stranger (n = 8; t_7_ = 0.695, p = 0.510; *Figure 2D*), demonstrating an impairment in pair bond formation. Conversely, control females had no partner preference after a 24-hour cohabitation (n = 6; t_5_ = 1.895, p = 0.117), although this lack of effect could be due to low sample size. However, defeated females had a significant preference for their partner over the stranger (n = 6; t_5_ = 8.268, p = 0.0004; *Figure 2E*), suggesting a potential promotion of pair bond formation as a result of defeat experience. In order to confirm this, the experiment was repeated with another cohort of females that cohabitated with a male for only 6 hours, which does not as reliably lead to partner preference [52]. Control females showed no preference for their partner over the stranger (n = 8; t_7_ = 0.801, p = 0.449) while defeated females again showed a significant partner preference (n = 9; t_8_ = 5.806, p = 0.0004; *Figure 2F*). These results support an impairment of pair bond formation as a result of defeat in males, but an unexpected defeat-induced acceleration of pair bond formation in females.

### Dual viral vector injection leads to selective expression in BNST CRHergic neurons

Because of the lack of CRH-Cre transgenic lines in prairie voles, selective chemogenetic manipulation of BNST CRHergic neurons was accomplished through an injection of a viral vector cocktail of CRH-Cre with Cre-inducible (DIO) mCherry (control), hM3DGq-mCherry (excitatory), or hM4DGi-mCherry (inhibitory) directly into the BNST (*Figure 3A*). First, we confirmed that the CRH-Cre vector led to selective expression in CRHergic neurons. A small cohort of prairie voles was injected with the CRH-Cre/DIO-mCherry cocktail into the BNST followed by an i.c.v. injection of colchicine 12 days later. There was significant overlap of CRH-ir cells with mCherry-expressing cells (mCherry cells co-labeled with green CRH-ir = 92.9% ± 2.3 %; *Figure 3B-C*). This confirmed that any effects of DREADD expression would be due to selective manipulation of BNST CRHergic neurons.

**Figure 3:**
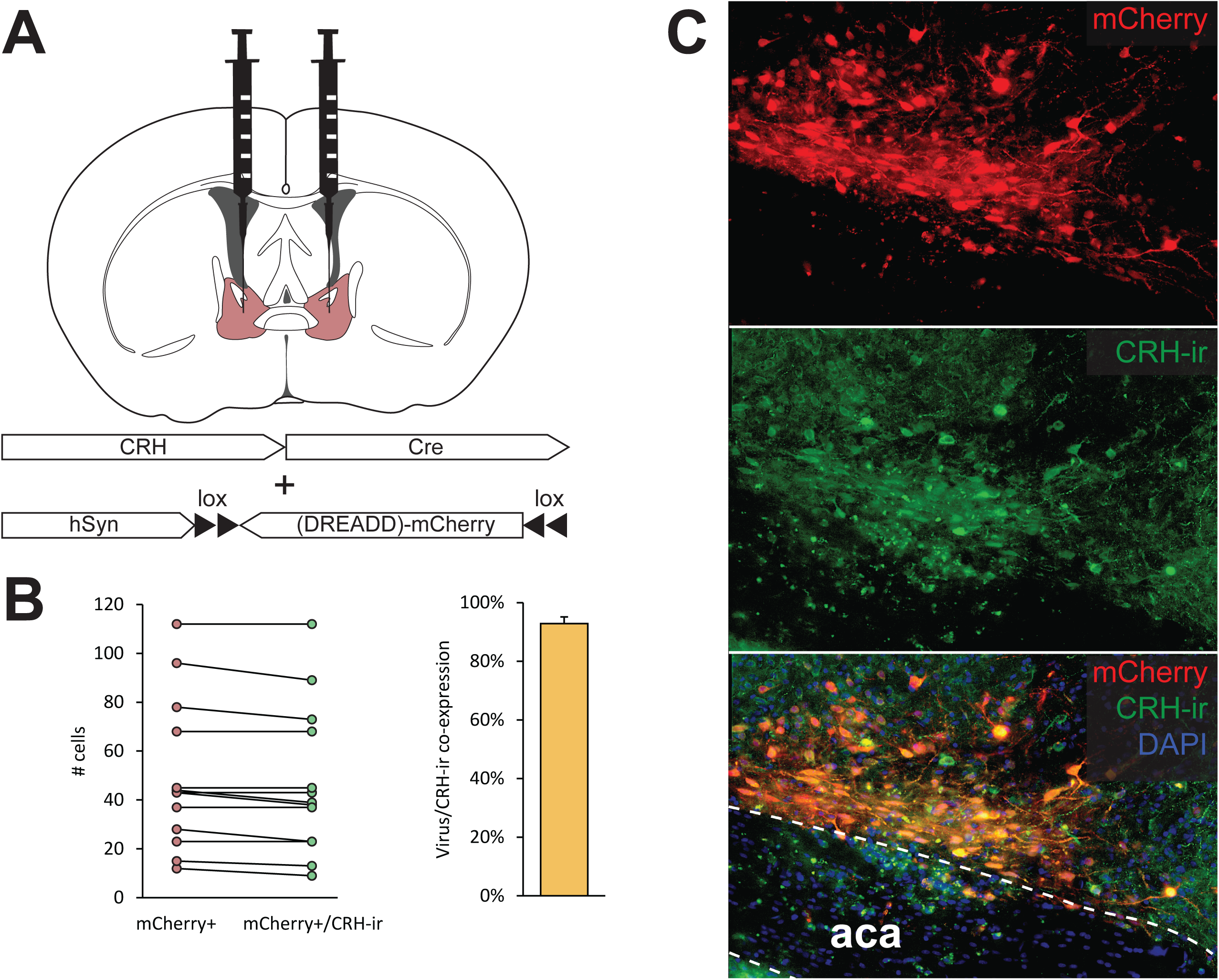
Experimental design for chemogenetic manipulation of BNST CRHergic neurons. A) Target site for bilateral viral injection and schematic for selective expression of DREADD protein and/or control mCherry in CRHergic neurons. B) Viral expression co-label with CRH immunoreactivity (CRH-ir). Validation of the dual-viral vector injection revealed 93±2% CRH-ir on mCherry-positive cells, indicating high selectivity. C) Representative image of selective viral expression in CRHergic neurons. Viral expression localized to the dorsal BNST.

### BNST CRHergic neurons have a stress-dependent role in pair bonding in male prairie voles

Next, the effect of BNST CRHergic activation on pair bond formation in male prairie voles was determined. Two weeks after the mCherry or Gq viral vector cocktail injection, male voles were injected with 3 mg/kg CNO and paired with a female for 6 hours, which is insufficient to lead to partner preference in male voles under normal conditions [56]. Female partners of male subjects in this experiment were not EB-primed to make any enhancing effects of BNST CRHergic stimulation on pair bond formation more apparent. Following the 6-hour cohabitation, the mCherry group indeed did not demonstrate a significant preference for their partner over the stranger (n = 8; t_7_ = 0.121, p = 0.907). However, the Gq group had a significant preference for their partner over the stranger after only 6 hours of cohabitation (n = 11; t_10_ = 2.232, p = 0.049; *Figure 4B*). This confirmed that activating BNST CRHergic neurons in males accelerates pair bond formation.

**Figure 4:**
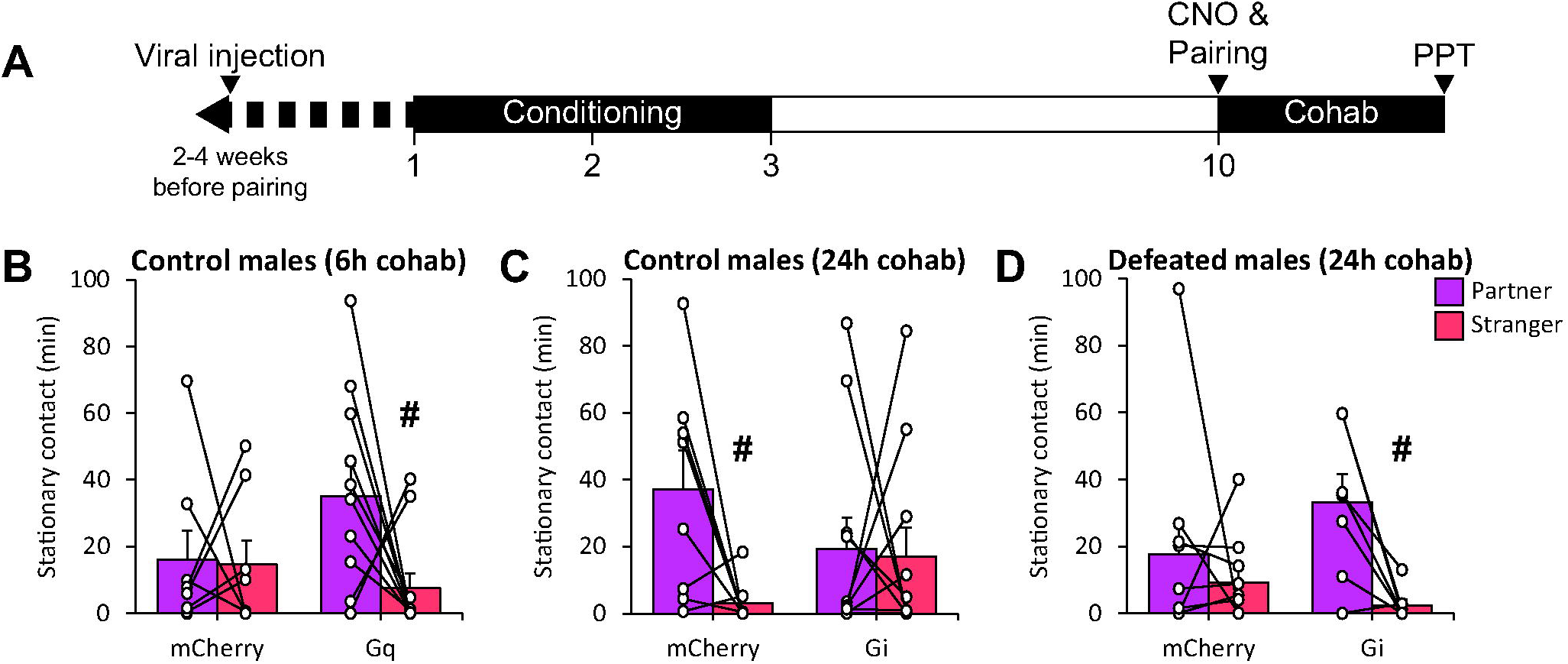
Effect of BNST CRHergic manipulation on partner preference in stress-naive and defeated male prairie voles. A) Experimental timeline. Injection of the CRH-Cre/Cre-inducible virus into the BNST occurred 2-4 weeks prior to pairing (1-3 weeks prior to conditioning). Animals were paired with a female partner for 6 or 24 hours then pair bond status was assessed with PPT. B) Activation of BNST CRHergic neurons with Gq-DREADD expression promoted partner preference in stress-naïve males (n = 8 mCherry/11 Gq). C) Inhibition of BNST CRHergic neurons with Gi-DREADD expression prevented partner preference in stress-naïve males (n = 8 mCherry/11 Gi). D) Inhibition of BNST CRHergic neurons with Gi-DREADD expression reversed defeat-induced impairments in pair bonding in male voles (n = 10 mCherry/7 Gi). #p < 0.05 vs. partner.

Next we aimed to determine whether eliminating normal BNST CRHergic signaling during cohabitation would prevent the formation of a pair bond. Males expressing mCherry or Gi were injected with CNO and cohabitated with a sexually receptive female partner for 24 hours. Following this cohabitation, mCherry males demonstrated the expected preference for their partner over the stranger female (n = 8; t_7_ = 2.626, p = 0.034). Conversely, Gi males had no preference between their partner and the stranger (n = 11; t_10_ = 0.155, p = 0.880; *Figure 4C*), confirming that BNST CRHergic neuronal activity is necessary for normal pair bond formation in males.

Earlier studies performed in mice have found that social defeat leads to an increase in CRH production in males [30], which may suggest that BNST CRH activity has a negative correlation with affiliation and bond formation. Yet we demonstrated earlier that pair bonding is also associated with an increase in BNST CRH production in stress-naïve animals. The BNST can differentially modulate social and approach/aversion behavior based on prior stress experience [20,33,57]. Thus, we hypothesized that the defeat experience would switch BNST CRHergic activity from promoting to preventing pair bond formation in males. Therefore, BNST CRHergic inhibition was repeated in defeated animals to see if decreasing CRHergic activity would reverse the effect of defeat on pair bond formation. In defeated males, mCherry-expressing animals had no preference between the partner and the stranger (n = 10; t_9_ = 0.740, p = 0.478). However, Gi-expressing males demonstrated a rescue of partner preference behavior after the 24-hour cohabitation (n = 7; t_6_ = 3.458, p = 0.014; *Figure 4D*).

### Stress uncouples BNST CRHergic neurons and pair bonding in female prairie voles

The above experiments were repeated in females. Following 6 hours of cohabitation, the non-defeated females in both the mCherry (n = 8; t_7_ = 3.939, p = 0.006) and Gq groups (n = 8; t_7_ = 3.348, p = 0.012) demonstrated a significant partner preference *(Figure 5B*). Because of the significant preference in the mCherry group, it is impossible to determine whether Gq DREADD activation of BNST CRHergic neurons is sufficient to induce partner preference as there could potentially be a ceiling effect. CNO has been demonstrated in the past to have potential off-target effects [58], so we confirmed that CNO alone was not leading to accelerated partner preference by treating a separate cohort of females with either vehicle or CNO prior to the 6-hour cohabitation with a male. Neither vehicle-treated (n = 6; t_5_ = 1.868, p = 0.121) nor CNO-treated females (n = 6; t_5_ = 0.173, p = 0.869) exhibited a partner preference following the 6-hour cohabitation (*Supplemental Figure 1*). Therefore, the sufficiency of BNST CRHergic neuronal activation to induce pair bond formation in female prairie voles is inconclusive.

**Figure 5:**
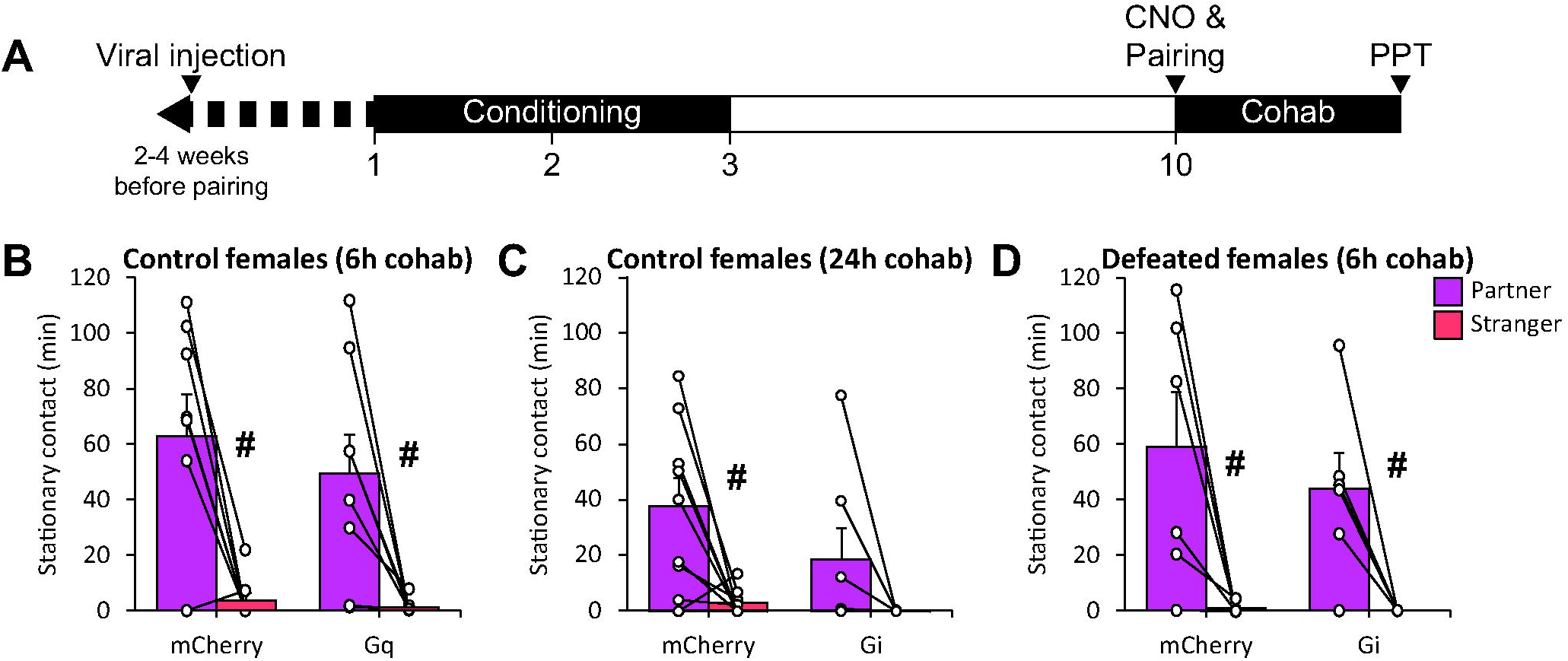
Effect of BNST CRHergic manipulation on partner preference in stress-naive and defeated female prairie voles. A) Experimental timeline. Injection of the CRH-Cre/Cre-inducible virus into the BNST occurred 2-4 weeks prior to pairing (1-3 weeks prior to conditioning). Animals were paired with a male partner for 6 or 24 hours then pair bond status was assessed with PPT. B) The effects of BNST CRHergic neuronal activation on pair bonding in stress-naïve females is unclear due to a potential ceiling effect (n = 8 mCherry/8 Gq). C) Inhibition of BNST CRHergic neurons with Gi-DREADD expression prevented pair bond formation in stress-naïve females (n = 9 mCherry/7 Gi). D) Inhibition of BNST CRHergic neurons with Gi-DREADD failed to prevent pair bond formation in defeated females (n = 6 mCherry/6 Gi). #p<0.05, ##p<0.01 vs partner.

Following the CNO injection and 24-hour cohabitation with a male partner, non-defeated mCherry females had a significant partner preference (n = 9; t_8_ = 3.230, p = 0.012). However, the Gi group had no significant preference between the partner and the stranger (n = 7; t_6_ = 1.649, p = 0.150; *Figure 5C*). This finding supports the necessity of this signaling in normal pair bond formation in females similar to that in males.

In defeated females, mCherry-expressing animals had a significant preference for their partner over the stranger (n = 6; t_5_ = 2.898, p = 0.034) after a 6-hour cohabitation, in line with what was demonstrated in the earlier experiment. However, following defeat experience, Gi inhibition of BNST CRHergic activity no longer prevented partner preference behavior (n = 6; t_5_ = 3.402, p = 0.019; *Figure 5D*), indicating defeat uncouples BNST CRH neurons from regulating pair bond formation in females.

## Discussion

In this study, we aimed to determine the effect of social defeat on pair bond formation in prairie voles of both sexes. Based on research revealing that a chronic stressor leads to impairments in pair bonding in male prairie voles [44], and our prior work demonstrating that there are no sex differences in social avoidance following defeat [39,40], we initially hypothesized that defeat experience would cause an impairment in pair bond formation in both sexes. However, while pair bonding was indeed impaired in defeated males, defeat unexpectedly led to an acceleration of pair bonding in females.

This is not the only instance of stress having opposing effects on pair bond behavior based on sex; an acute stressor has been shown to promote partner preference in male voles and impair partner preference in female voles [59]. Notably, this is the opposite effect from what was observed in the present study. This difference may largely be due to the effects of acute vs. chronic stress, as chronic stress causes an impairment in pair bond formation in males [44]. Additionally, the social nature of social defeat is likely a significant contributor to the impact on pair bond formation, since the social rejection experienced during social defeat is widely generalized to all unknown conspecifics [39,40,42]. We demonstrate here that this social avoidance is even generalized to potential mates, meaning that the social withdrawal state induced by social defeat experience cannot be overcome even by sexual motivation.

We also demonstrate here for the first time a novel neurobiological mechanism of pair bond formation with the activation of BNST CRHergic neurons. The connection between BNST CRHergic neurons and pair bond formation has been suggested by prior literature [38,60], but a causative relationship was not confirmed. Through selective chemogenetic manipulation of BNST CRHergic neurons, we were able to confirm the necessity of BNST CRHergic neuronal activity for pair bonding in prairie voles of both sexes. While we were able to also confirm the sufficiency of BNST CRH neuronal activity to induce pair bond formation in males naïve to stress, the results of BNST CRHergic activation in stress-naïve females is unfortunately inconclusive. The 6- and 24-hour cohabitation periods are standard in the prairie vole field for being respectively “insufficient” and “sufficient” for pair bond formation in both sexes [46,51,52,56]. However, there are studies that show similar inconsistency in the 6-hour cohabitation time with females in that some studies show indication of pair bond formation in females with this shorter cohabitation time [53]. We did confirm that off-target effects of CNO were not contributing to this inconsistency in females; thus, further experimentation with perhaps an even further truncated cohabitation time may be necessary to determine whether activation of BNST CRH neurons is sufficient to accelerate pair bond formation in stress-naïve females. Despite this limitation, we have confirmed within this study that BNST CRHergic activity is necessary for pair bond formation in both sexes, and sufficient to induce it in at least males. The causational positive relationship between BNST CRHergic neuronal activity and pair bond formation demonstrated here stands in stark contrast to the typical relationship between BNST CRHergic activation and negative valence. While knockout of CRH in GABAergic forebrain neurons does diminish social interaction [33], this knockout encompasses CRH neurons in both the BNST and central amygdala, which have been demonstrated to have opposing roles in promoting motivation [61,62]. Despite this, we show here that activation of BNST CRHergic neurons is both sufficient and necessary for the promotion of pair bond formation, until a social stressor is experienced, indicating that the role of BNST CRHergic neurons in motivational valence is not as simple as promoting approach or aversion and may be dependent on social context.

It is interesting to note then that only lower doses of centrally-administered CRH promote partner preference, and higher doses do not [34]. Social defeat leads to an increase in BNST CRH production in male mice [30], and this was also observed in defeated male prairie voles. Thus, it is possible that an overabundance of CRH in the BNST following social defeat contributes to the stress-dependent role of BNST CRHergic neurons in pair bonding in male prairie voles. The stress-dependent role of the BNST in behavior is a well-documented phenomenon that significantly contributes to the putative role of the BNST in “valence surveillance” [20,33,57]. This is likely modulated by BNST projections to the ventral tegmental area (VTA), which can promote either an anxiogenic or anxiolytic state depending on the proportion of glutamatergic to GABAergic signaling, respectively [19,63]. Long-range GABAergic CRH neurons projecting from the BNST to the VTA contribute directly to this anxiolytic effect, as knockout of CRH from these neurons promotes anxiety-like behavior and negatively impacts dopamine release from the VTA [33]. The majority of CRHergic neurons within the BNST are GABAergic [64], and although long-range GABAergic projections underlie the established connection with the VTA, there are also CRH-GABA interneurons within the BNST that likely modulate these long-range projections [65]. Interestingly, we also found here that both selective activation and selective inhibition of BNST CRHergic neurons leads to an overall increase in BNST CRHergic activity. With the calcium indicator expressing non-selectively in the BNST, it is possible that this overall increase in activity upon chemogenetic manipulation of CRHergic neurons reflects a shift in cell activity dynamics depending on CRH activity.

Such a shift in cell activity dynamics likely occur naturally following stress, as chronic stress leads to increased connectivity between CRHergic neurons within the BNST, leading to stress-dependent reciprocal inhibition of CRHergic neurons [65]. Thus, the potential overabundance of CRH within the BNST following defeat and cohabitation may lead to higher CRH-CRH connectivity within the BNST, inhibiting BNST CRH long-range signaling and disinhibiting VTA GABAergic signaling, subsequently leading to an anxiogenic state [66]. Alternatively, BNST CRHergic projections to the nucleus accumbens (NAc) may contribute to this stress-dependent regulation of pair bond formation as well. Although a BNST^CRH^➜NAc pathway is not as well established as the BNST^CRH^➜VTA pathway, there is evidence in male mice and rats that such a projection does exist [62,67,68], and conserved in humans [69]. This signaling pathway is potentially incredibly relevant to the stress-induced alterations in pair bond formation presented in the current study; CRH within the NAc not only promotes pair bond formation in stress-naïve male prairie voles [70], but also switches from an appetitive to an aversive signal following stress experience in mice through modulation of dopamine signaling [71]. With mesolimbic dopamine being a major neurobiological mechanism of pair bond formation [72], this proposed circuit of stress-induced anxiogenic shift in BNST^CRH^➜VTA and/or BNST^CRH^➜NAc signaling certainly warrants further investigation in the examination of the neurobiological connection between stress and affiliation.

To our knowledge the present study is the first characterization of the effects of a chronic stress state on pair bonding in female prairie voles. Perhaps one of the most striking findings was the discovery of the sexually dimorphic effect of defeat on pair bond formation. With no sex differences in social investigation following social defeat [39,40], this effect was unexpected. Furthermore, the stress-dependent role of BNST CRHergic neurons in pair bonding also differed between males and females. Defeat increased BNST CRH mRNA in males but not females while establishing a male-female pair increased BNST CRH mRNA expression in females but not males. This is not necessarily surprising, as the BNST and CRHergic neurons within it are well established to be sexually dimorphic [73–75]. Despite this, research investigating the role of the BNST in “valence surveillance” and the stress-dependent anxiogenic/anxiolytic shift in this region has been almost exclusively performed in males. Research in California mice has shown that oxytocinergic signaling within the BNST contributes to sex differences in social approach and vigilance [76]. Notably, CRH and oxytocin have been demonstrated in rats to have a bidirectional modulatory relationship with each other [64,77,78]. Oxytocin signaling modulates the behavioral response to social defeat in both prairie voles and Mandarin voles [39,42,79], and is a well-established regulator of pair bond formation in prairie voles [80,81]. Thus, while the lack of research investigating the effects of stress on the BNST in females makes it difficult to speculate how BNST CRH signaling could become uncoupled from pair bond regulation as a result of defeat, bidirectional regulation between oxytocin and CRH could be a compensatory mechanism promoting pair bond formation following stress experience in females.

While the role of BNST CRHergic signaling in modulating social behavior and affiliation would be difficult to assess in humans, general BNST activity is known to be impacted in individuals suffering from social anxiety disorder [17,82]. Much like animal models, unfortunately, the role of the BNST in social interaction and affiliation is not as thoroughly characterized as its role in stress and threat anticipation in humans. The results of the present study place BNST CRHergic neurons at the intersection of the role of the BNST in both stress and affiliation. Further investigation of this mechanism could provide new insight on how to approach managing deficits in social connections in those suffering from social anxiety or other conditions that result from severe social stressors.

## Author contributions

MT contributed to conceptualization, formal analysis, investigation, methodology, validation, visualization, writing of the original draft, and review and editing of the manuscript. JG, EV, and LN contributed to formal analysis, investigation, validation, and review and editing of the manuscript. AS contributed to conceptualization, formal analysis, funding acquisition, methodology, project administration, resources, supervision, validation, and review and editing of the manuscript.

## Acknowledgements

We thank K. Gossman for his assistance with validating specificity of viral expression with immunofluorescence, and the KU Animal Care Unit staff for their care of the prairie vole colonies. Research reported in this publication was supported by the National Institute of Neurological Disorders and Stroke of the National Institutes of Health under Award Number R01NS113104. The content is solely the responsibility of the authors and does not necessarily represent the official views of the National Institutes of Health. Additional funding was provided by K-INBRE.

