## Supplemental Figure 1 for "Extended amygdala corticotropin-releasing hormone neurons regulate sexually dimorphic changes in pair bond formation following social defeat in prairie voles (*Microtus ochrogaster*)"

Supplemental Figure 1: CNO administration does not significantly promote partner preference in female voles. Female prairie voles received a sham injection into the BNST, then two weeks later were administered either vehicle (n=5) or 3 mg/kg CNO (n=6) prior to a 6-hour cohabitation with a male partner. Neither group had a significant preference for the partner over the stranger.

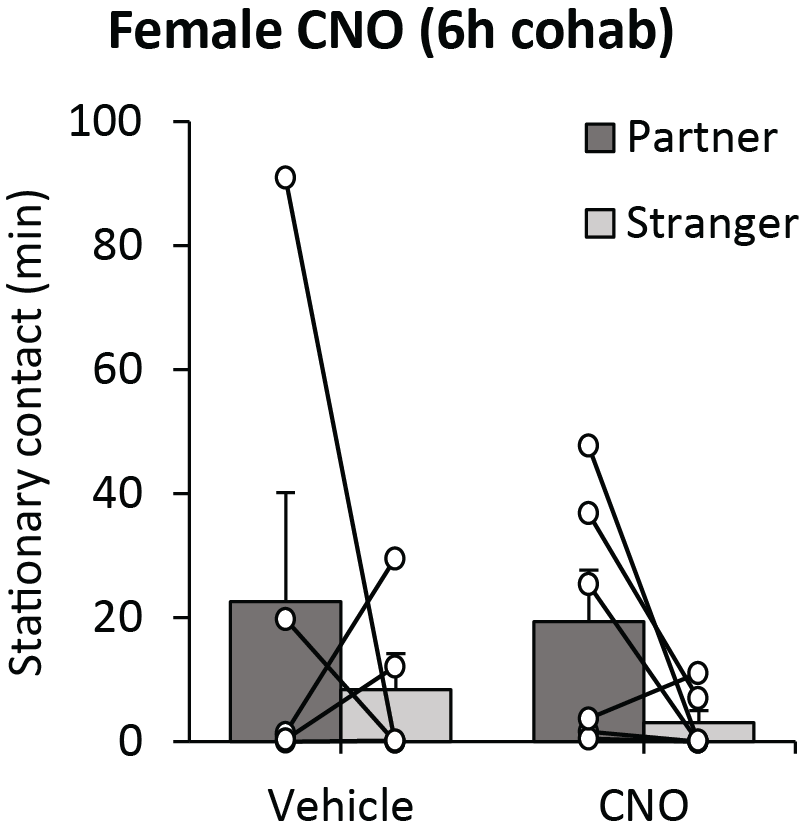

**Key resources table**

| REAGENT or RESOURCE | SOURCE | IDENTIFIER |
| --- | --- | --- |
| Antibodies | | |
| Guinea pig anti-CRH | Peninsula Laboratories International | Cat #T-5007; RRID: AB_518256 |
| Goat anti-guinea pig IgG (Alexa Fluor 488) | Invitrogen | Cat #A11073; RRID: AB_2534117 |
| Bacterial and virus strains | | |
| AAV2-CRH-Cre | Biohippo | Cat #PT-0588 |
| AAV2-hSyn-DIO-mCherry | Bryan Roth (Addgene) | Cat #50459; RRID: Addgene_50459 |
| AAV-hSyn-DIO-hM3D(Gq)-mCherry | Bryan Roth (Addgene) | Cat #44361; RRID: Addgene_44361 |
| AAV2-hSyn-DIO-hM4D(Gi)-mCherry | Bryan Roth (Addgene) | Cat #44362; RRID: Addgene_44362 |
| Chemicals, peptides, and recombinant proteins | | |
| Colchicine | Thermo Scientific | Cat #J61072ME |
| Clozapine-N-Oxide | NIMH Chemical Synthesis and Drug Supply Program | N/A |
| Critical commercial assays | | |
| BCA assay | Thermo Scientific | Cat #23246 |
| RNEasy Mini Kit | Qiagen | Cat #74104 |
| Qubit RNA High Sensitivity kit | Invitrogen | Cat #Q32852 |
| Reverse transcription kit | Applied Biosystems | Cat #4368814 |
| Experimental models: Organisms/strains | | |
| *M. ochrogaster* | Breeding colony | N/A |
| Oligonucleotides | | |
| CRH forward primer;  5’ – GGG GAA CCT CAA CAG AAG CC – 3’ | IDT | N/A |
| CRH reverse primer;  5’ – ACA CGC GGA AAA AGT TAG CC – 3’ | IDT | N/A |
| HPRT forward primer;  5’ – CCC AGC GTC GTG ATT AGT GA – 3’ | IDT | N/A |
| HPRT reverse primer;  5’ – TCG AGC CAG TCT TTC AGT CC – 3’ | IDT | N/A |
| Software and algorithms | | |
| ImageJ | Schneider et al.^57^ | https://imagej.nih.gov/ij/ |
| JWatcher | UCLA | https://www.jwatcher.ucla.edu/ |
